# Body size, not species identity, drives body heating in alpine *Erebia* butterflies

**DOI:** 10.1101/2022.10.03.510594

**Authors:** Irena Kleckova, Jan Okrouhlik, Tomas Svozil, Pável Matos-Maraví, Jan Klecka

## Abstract

Efficient thermoregulation is crucial for animals living under fluctuating climatic and weather conditions. We studied the body heating of six butterfly species of the genus *Erebia* (Lepidoptera: Nymphalidae) that co-occur in the European Alps. We tested whether butterfly physical characteristics (body size, wing loading) are responsible for the inter-specific differences in body temperatures recorded previously under natural conditions. We used a thermal camera to measure body heating of wild butterfly individuals in a laboratory experiment with artificial light and heating sources. We revealed that physical characteristics had a small effect on explaining inter-specific differences in mean body temperatures recorded in the field. Our results show that larger butterflies, with higher weight and wing loading, heated up more slowly but reached the same asymptotic body temperature as smaller butterflies. Altogether, our results suggest that differences in body temperatures among *Erebia* species observed in the field might be caused mainly by species-specific microhabitat use and point towards an important role of active behavioural thermoregulation in adult butterflies. We speculate that microclimate heterogeneity in mountain habitats facilitates behavioural thermoregulation of adults. Similarly, microclimate structuring might also increase survival of less mobile butterfly life stages, i.e., eggs, larvae and pupae. Thus, landscape heterogeneity in management practices may facilitate long term survival of montane invertebrates under increased anthropogenic pressures.

## 1. Introduction

Thermal ecophysiology of butterflies is driven by interactions between the environment and species-specific physical (morphology, physiology) and behavioural thermoregulation. Important morphological traits affecting butterfly body temperatures include body size (Heinrich, 1986; Kleckova et al., 2014), wing loading (i.e., the ratio of body weight over wing area) and colouration (Kingsolver, 1983). Behaviourally, insects thermoregulate by specific body postures (Tsai et al., 2020) or by microhabitat choice (Kleckova & Klecka, 2016). Adult butterflies increase their body temperature over the ambient air temperature by basking postures (Heinrich, 1995; Kingsolver, 1985a) and by conduction from warm stones or soil (Kemp & Krockenberger, 2002; Kingsolver, 1985b; Kleckova et al., 2014). Conversely, body temperature can be decreased by a shift to a cooler microhabitat (Dreisig, 1995; Shreeve, 1984; Wickman, 1988). Hence, body temperatures in wild butterflies are likely driven by the interaction of morphological and physical traits with active behavioural thermoregulation.

However, our knowledge about the interactions between behavioural thermoregulation and physical differences is scarce (Abram et al., 2017) because they are difficult to disentangle by field observations (e.g. Lawson,et al., 2014a). Adult butterflies likely respond to short-term environmental changes by active thermoregulation based on their species-specific physiological optima (Bladon et al., 2020; Kleckova et al., 2014; Kleckova & Klecka, 2016). The physiological optima are, in turn, also affected by the active choice of ambient temperature and daily activity patterns of different species of butterflies (Kingsolver, 1983) and other insects, such as beetles (Gallego et al., 2018). The interplay among behaviour, morphology and physiology on thermoregulatory strategies has been observed also at the inter-population level in butterflies (Nève & Després, 2020) and intra-population level in grasshoppers (Forsman, 2000). Further, thermoregulatory behaviour of butterflies is tightly connected with morphology as behaviour prevents, for example, overheating of living cells distributed in the wings veins (Tsai et al., 2020). We still know little about the main traits involved in thermoregulation of butterflies occurring in extreme environments such as alpine regions.

To respond to changes of local climatic conditions in mountains (Rödder et al., 2021), butterflies may exploit different local physiological optima (MacLean et al., 2016), might disperse to optimal climates (Konvicka et al., 2003) or change their thermoregulatory behaviour, e.g., by shifts in the timing of daily activities (Cros et al., 1997; Slamova et al., 2011) and microhabitat use (Kirkpatrick & Sheldon, 2022; Wilson et al., 2015). Further, climate change drives the timing of seasonal activities, mainly by affecting less mobile stages of butterfly life cycle (Konvicka et al., 2021; Radchuk et al., 2013). Species, populations and even individuals differ in their responses to ambient temperatures. Such selection pressures of local conditions probably resulted in variable body temperatures necessary for flight initiation in different populations of *Melitaea cinxia* butterflies (Advani et al., 2019). In addition, Matilla et al. (2015) detected differences in body temperature in flight among genotypes and sexes of *M. cinxia*.

To test the role of physical differences for body heating, we studied adult *Erebia* (Nymphalidae, Satyrinae) butterflies occurring in sympatry in alpine environments. Mountains provide high heterogeneity of thermal environments for thermoregulation of butterflies (Wickman, 2009). However, microclimatic conditions are changing rapidly (Stuhldreher & Fartmann, 2018) and many alpine species, including *Erebia*, are threatened by climate warming (Konvicka et al., 2021; Sistri et al., 2022). *Erebia* butterflies have low mobility (Polic et al., 2014; Slamova et al., 2013), which probably leads to slower uphill shifts in response to climate warming compared to mobile generalist species (Rödder et al., 2021). On the other hand, at least some populations of high-altitude *Erebia* specialists use suitable habitats at altitudes below their climatic optimum (Cizek et al., 2003). Inter-specific differences in mean body temperatures of co-occurring closely related *Erebia* species might have been driven mainly by microhabitat use and body size (Kleckova et al., 2014), but it is unclear what the main thermoregulation strategy is. To test the relative effects of species identity (a proxy for possible physiological differences) and morphology (individual weight and wing-loading as drivers of passive heating), we measured body temperature heating rates under an artificial source of radiation. Our hypothesis was that different thermal conditions and microhabitats along elevational gradients might have driven divergent physiological constraints and heating rates may differ among species. Further, we expected that larger individuals and individuals with lower wing loading would heat up more slowly than smaller individuals (Wickman, 2009), as detected in other butterflies such as *Pararge aegeria* (Berwaerts et al., 2001) and *Hypolimnas bolina* (Kemp & Krockenberger, 2004). Finally, by comparing data from the laboratory heating experiments to field measurements of body temperatures (Kleckova et al., 2014), we inferred the relative importance of microhabitat choice in determining mean adult body temperatures.

## 2. Material and methods

### 2.1. Study site and species

We collected adult butterflies of six species of *Erebia* in the Austrian Alps, close to the town of Sölden, Tirol (1900-2020 m a.s.l., 46°57’37”N 11°01’49”E) and in Hochgurgl (2150 m a.s.l., 46°54’6”N, 11°03’14”E, 7 km away). We collected 15-20 individuals of each species in the field, a subset was used to measure heating rates in the lab (Table 1). The additional specimens were used for trial measurements and kept as a backup for unexpected events such as escape or mortality. We collected undamaged butterflies with minimal wing-wear and scored their sex. *Erebia* species are protandric (Kuras et al., 2003), and because we were able to access the study location for a limited period of time in the beginning of the species’ flight period, we collected mainly males. A balanced number of males and females for the experiments was available only for *E. pandrose* (Table 1).

**Table 1.**
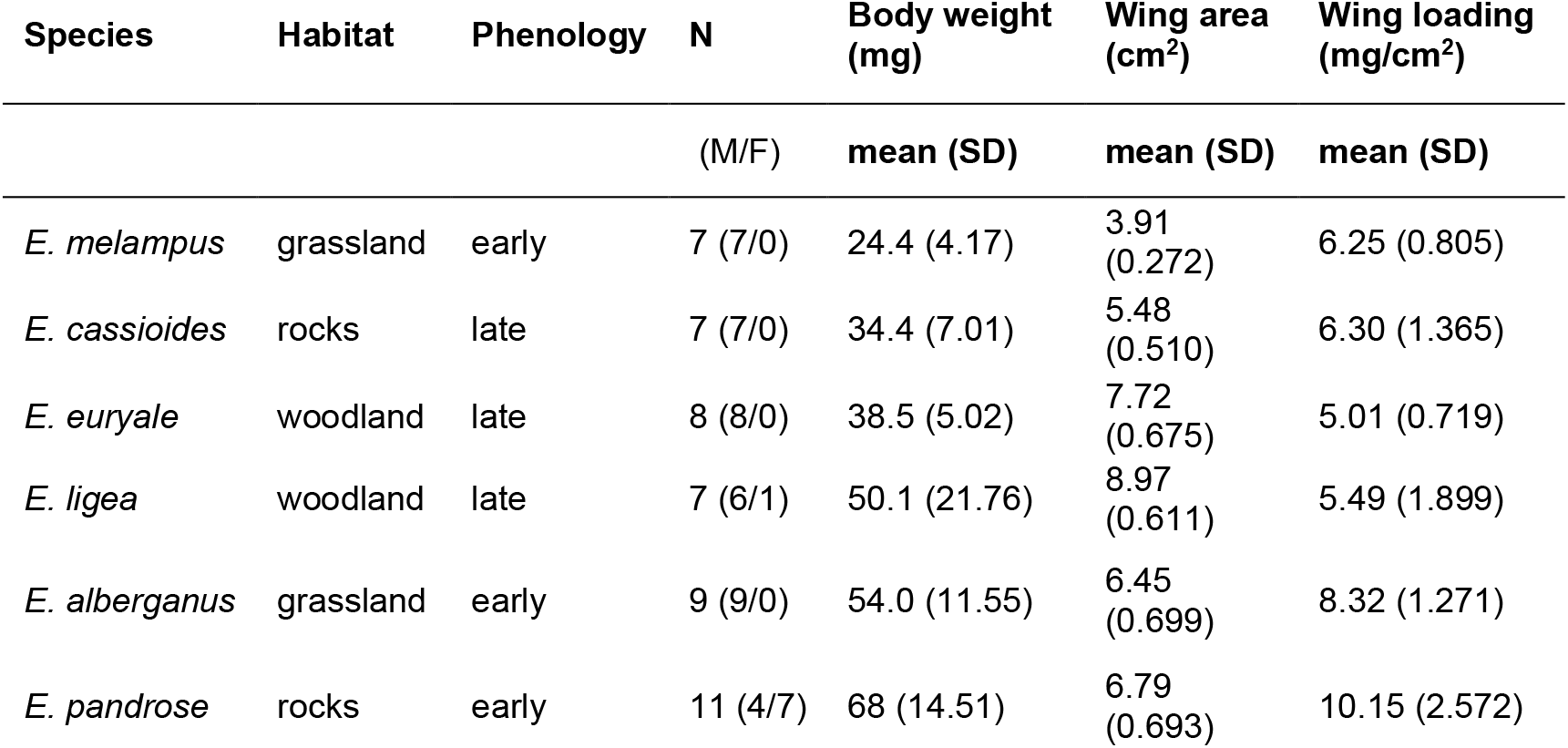
The list of study species of the butterfly genus *Erebia*. For each species, we provide information about its typical habitat, phenology (adults flying early or late in the summer), the number of individuals used for the lab measurements (N, males (M)/females (F)), their mean body weight, total wing area, and wing loading (body weight/wing area). Species are sorted according to their body weight from lightest to heaviest.

Species differ in their habitat preferences, altitude of occurrence, body size, and phenology. Rocky habitats above the treeline host *E. pandrose* (Borkhausen, 1788) and *E. cassioides* (Reiner & Hochenwarth, 1792), which increase their body temperature in warm stony microhabitats (Kleckova et al., 2014). Sub-alpine and alpine grassy meadows above the treeline are inhabited by *E. alberganus* (de Prunner, 1798) and *E. melampus* (Fuessly, 1775), which typically bask on stems of grasses. Subalpine meadows around the treeline and woodland clearings below the treeline are inhabited by *E. euryale* (Esper, 1805) and *E. ligea* (Linnaeus, 1758). These species bask on grass blades and on tree branches. All species have wings with dark brown upper side with orange spots. *E. pandrose* and *E. cassioides* have silver underside and higher proportion of orange colouration. The remaining four species have brownish underside. Regarding size differences, *E. cassioides* and *E. melampus* are small (forewing length less than 20 mm), whereas the other four species are large (forewing length above 20 mm). Three early flying species, *E. alberganus, E. melampus* and *E. pandrose*, were caught between 27^th^ and 29^th^ June 2011. Three late flying species, *E. ligea, E. cassioides* and *E. euryale*, were caught between 2^nd^ and 4^th^ August 2011. The butterflies were collected by hand netting and stored alive in envelopes inside a cooling box with suitable humidity provided by wet tissues (temperature range in the box was 10-15°C). We transported the specimens to the facilities of the Faculty of Science, University of South Bohemia in České Budějovice, Czech Republic, for the laboratory experiments.

### 2.2. Laboratory measurements of body heating

Before the experiments, the butterflies were acclimatised overnight, for at least 15 hours, in a net cage in a climate-controlled room (20°C) with 16 h day and 8 h night photoperiod. Butterflies were fed ad libitum by 20% sugar solution applied on the net. We measured the heating rate using a thermal camera (FLIR P660, Flir Systems) in a climate-controlled room. The thermal camera was mounted on a tripod 0.60 m above a flat wooden arena where the butterfly was placed. We used wood as material for the arena because it kept a relatively stable surface temperature during the experiments. On the camera, we set-up the emissivity to 0.85, humidity to 66 %, and object distance to 0.60 m for all individual specimens. The emissivity value compensates for the reflection, transmission, and absorption of IR energy and corresponds to a conservative estimate for insect cuticle, e.g. Verdu et al. (2012). The humidity value corresponds to the average air humidity in the experimental room (range 56-82%).

The acclimatised butterflies were first cooled at 5 °C for 10 minutes in a fridge with the lights on to keep the butterflies active (acclimatization in the dark induced resting behaviour and butterflies did not stand but fell to one side when placed inside the arena). Then, using cold forceps, they were placed in the middle of the experimental arena under two halogen light bulbs (18 W and 160 lm, irradiance at butterfly distance was 45 W/m^2^) situated 25 cm above the arena. However, butterflies were allowed to move across thermally homogeneous arena and they could change their body posture and open their wings. The butterfly surface body temperature *T*_*b*_ was recorded every 20 seconds for 10 minutes (Figure 1). In total, we measured the heating rate in 58 individuals. Nine individuals which escaped from the arena by “jumping” behaviour sooner than in two minutes (corresponding to 5 thermal camera images) were removed from subsequent analyses. The remaining 49 individuals were sitting during the experiment until they took off. Each individual was exposed to the experimental warming only once. The temperature in a middle point on the mesothorax was recorded using FLIR QuickReport©1.2 software, using a single manually selected point. The air-conditioning system in the experimental room created a strong draft, so we kept it turned off for the duration of the measurements. Consequently, the temperature in the experimental room fluctuated from 17.5 to 21.1 °C. To account for these fluctuations in the statistical analysis, room temperature (*T*_*room*_) and *humidity* was measured just before each trial.

**Figure 1.**
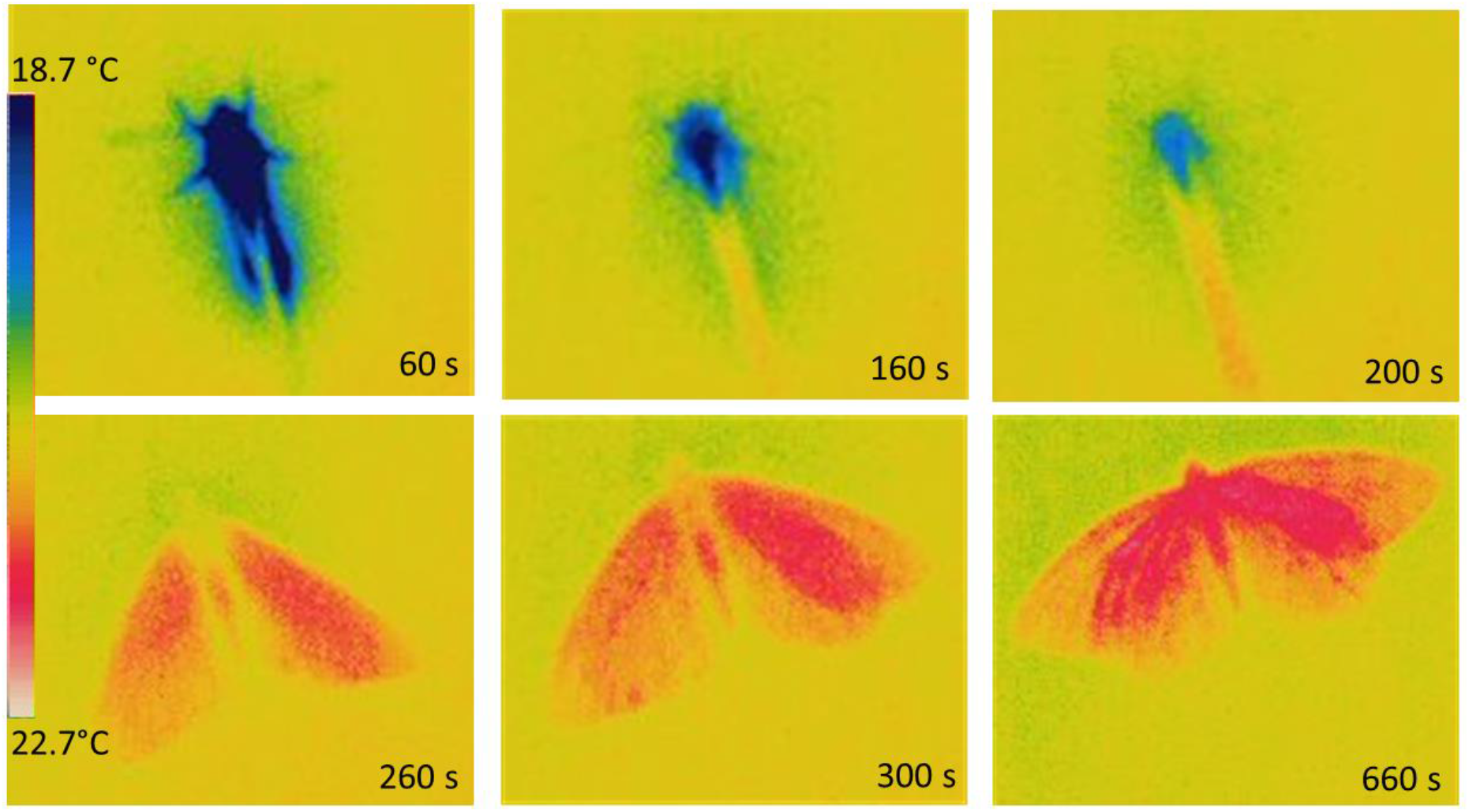
An example image from the thermal camera displaying the heating of an individual *Erebia* butterfly in *time* (seconds, s), blue indicates colder, red warmer temperatures. Typically, the butterflies were sitting initially with closed wings and open them later.

After the thermal camera measurements, the individual butterflies were sacrificed using ethyl-acetate. We obtained the weight (mg) of fresh individuals using a laboratory balance (precision 0.0001 g). We also photographed the left side of wings of every individual with wings separated from the body. Photographs of the pair of the wings with a scale were taken by the Panasonic FZ5 Camera from the distance of 100 cm using focal length 432 mm. Then, we calculated the wing area in the software ImageJ v.1.46 (Rasband, 1997) and multiplied by two to obtain an estimate of the total wing area. We calculated wing loading (mg/cm^2^) of every individual as the ratio of body weight (mg) over wing area (cm^2^) (Table 1).

### 2.3 Statistical analyses

#### Heating curves and species effects

All analyses and data visualisation were done in R version 4.2.0 (R Core Team, 2022). First, we analysed heating curves, describing the increase of body temperature *T*_*b*_ in time during basking. The curves were analysed using nonlinear mixed effects models (NLME) with Gaussian error distribution in the nlme package for R (Pinheiro et al., 2022). The heating curve was estimated by two alternative models:

1. following the modified asymptotic exponential equation *T*_*b*_ = *T*_*s*_ (1–e^−*b* (*time*-*c*)^), where *T*_*s*_ is the estimated asymptotic temperature reached by an individual. The slope of the asymptotic exponential curve (*b*) describes the heating rate of the body temperature increase in time until it reaches asymptotic body temperature *T*_*s*_.
2. following the modified Michaelis-Menten equation *T*_*b*_ *= T*_*s*_*((time-c)/(time+K-c))*, where the parameter *T*_*s*_ is the same as above and *K* is analogous to the Michaelis constant in classic Michaelis-Menten equation, but here *K+c* is the time when *T*_*b*_ reaches 50 % of the final *T*_*s*_.

We modified the usual form of the exponential equation and the Michaelis-Menten equation to include the parameter *c* which accounts for the fact that the initial *T*_*b*_ at *time=*0 was >0 °C (Figure 2). Specifically, *c* is the estimate of how long it would take for the butterfly to increase its body temperature from *T*_*b*_=0 °C to the real initial *T*_*b*_ (which varied among individuals) under the experimental conditions. An exponential model (Umbers et al., 2013, Amore et al., 2017) as well as the Michaelis-Menten equation (Muñoz et al., 2005) were previously used to describe ectotherm heating, but we are not aware of similar studies comparing the fit of multiple models.

**Figure 2.**
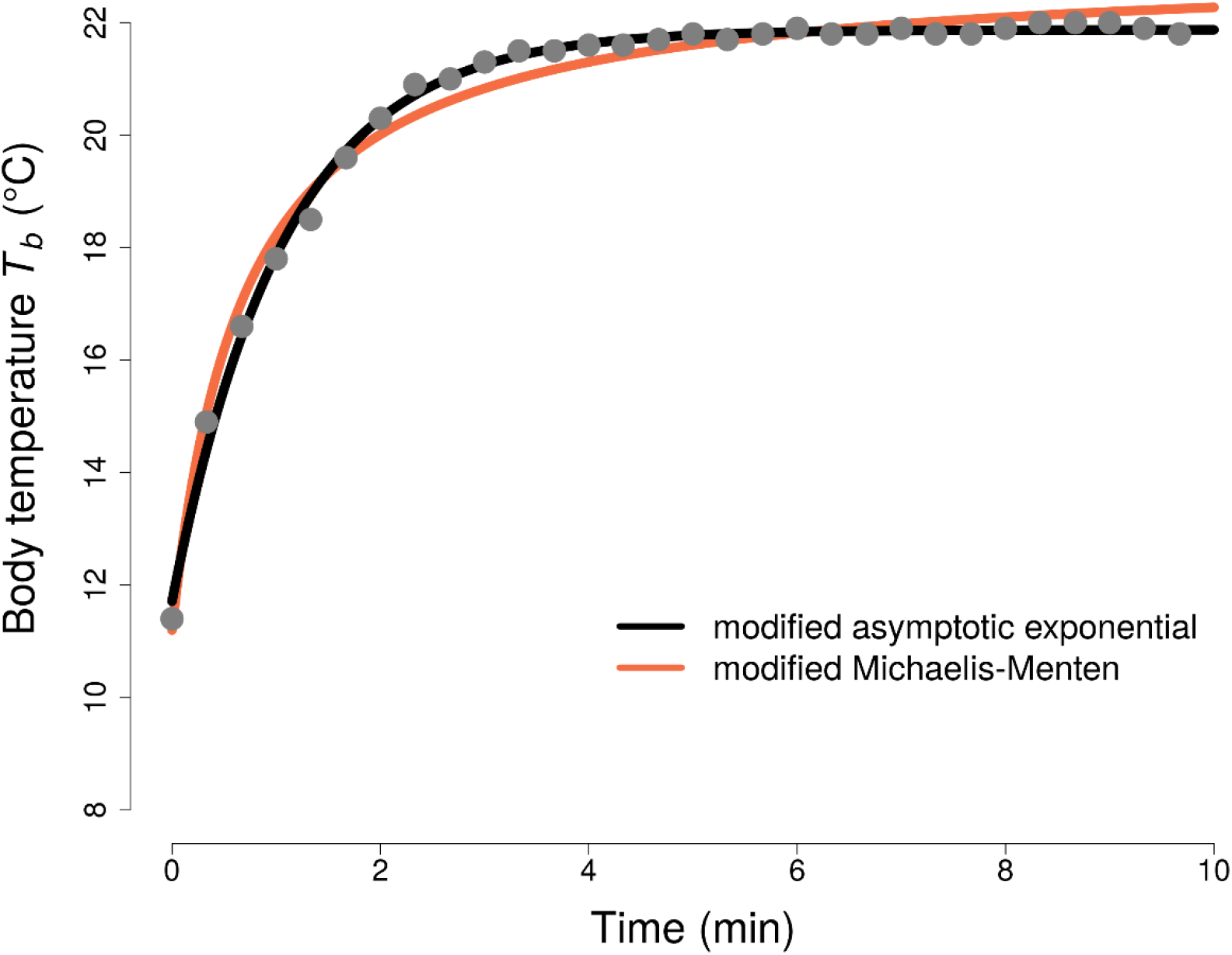
An example heating curve of a single individual of *Erebia melampus* butterfly with the comparison of the fit of the modified asymptotic exponential (black line) and Michaelis-Menten models (orange line). The modified asymptotic exponential model fits the data for this individual significantly better (ΔAIC = 38.0).

We compared the fit of the two models without any random effects and with two alternative random effect structures: the random effect of species and the random effect of individual nested within species. The individual identity was used to account for possible differences of the heating rate (*b*) and asymptotic body temperature (*T*_*s*_) in different individuals. The model fits were compared by the Akaike information criterion (AIC).

#### Body weight and wing loading effects

We tested how the shape of the heating curves depends on individual body mass and wing loading. We used generalised linear mixed effect models (GLMM) in the lme4 (Bates et al., 2015) and lmerTest libraries for R (Kuznetsova et al., 2017). For every individual, we extracted the estimates of parameters of asymptotic temperature *T*_*s*_ and heating rate *b* from the best fitting NLME model (i.e., modified asymptotic exponential, see Table 2). Then, we correlated the dependence of individual-level *T*_*s*_ or *b* on individual body weight and wing-loading. As additional predictors in the model, we included the *T*_*room*_ temperature and *humidity* in the experimental room recorded during individual measurements. We also compared mean body weight and wing-loading among species using generalised linear models (GLM) with Gamma distribution and log link function.

**Table 2.**
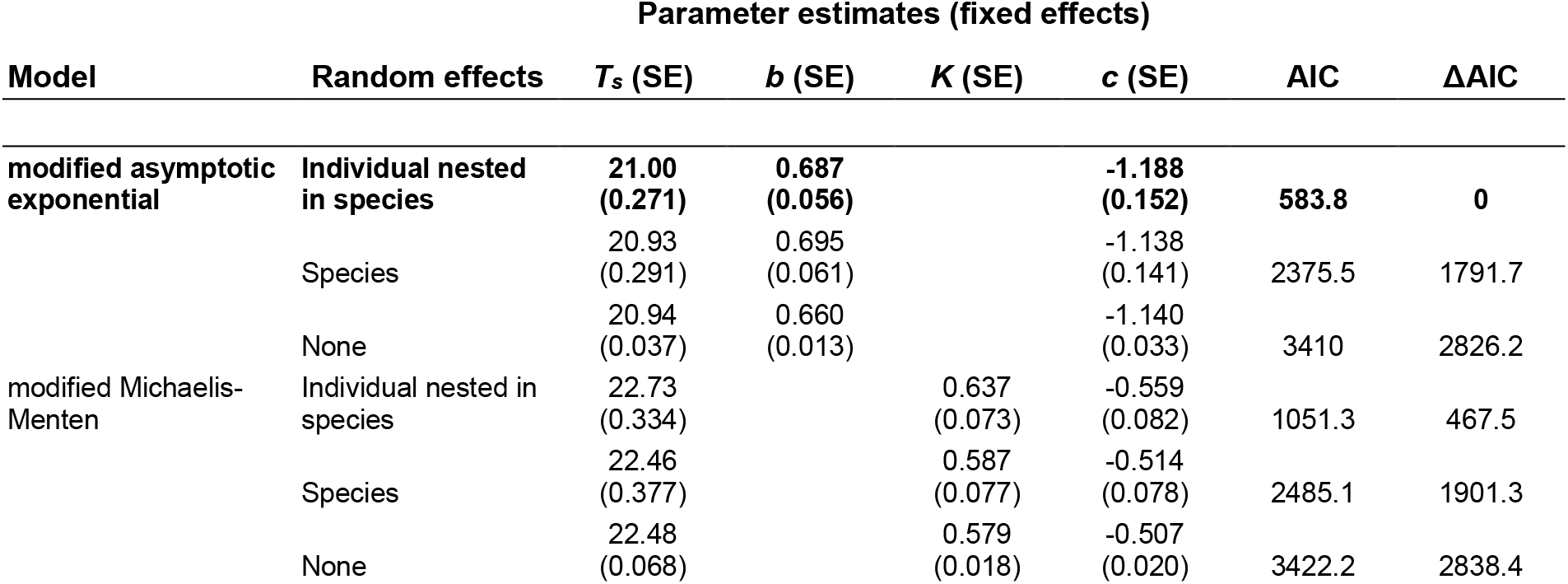
The comparison of the fit of the modified asymptotic exponential and Michaelis-Menten models describing butterfly heating under laboratory conditions. The models describe the increase of body temperature (*T*_*b*_) in *time* in 49 individuals of *Erebia* butterflies, which heated up under an artificial source of heating and radiation. Their *T*_*b*_ was measured by thermal camera. The non-linear mixed effect models included the random effects of individual nested in species and the random effect of species. The best model is highlighted **in bold**. Parameters describe maximum body temperature reached during heating (asymptotic temperature *T*_*s*_), heating rate (*b)*, Michaelis-Menten constant (*K*), and the estimated time it would take for the butterfly to increase its body temperature from *T*_*b*_=0 °C to the actual initial *T*_*b*_ (c).

#### Heating in the laboratory vs. body temperature in the field

We intended to compare species-specific values of the asymptotic temperature (*T*_*s*_) reached in laboratory conditions with body temperatures (*T*_*b_field*_) reached in the field (Kleckova et al., 2014). Because the range of ambient temperatures experienced in the laboratory differed from the range in the field, we estimated standardised asymptotic body temperature *T*_*s*_ and standardised body temperature in the field *T*_*b_field*_. In the field, the experienced air and temperature ranged from 12.0 to 25.0 °C and microhabitat temperature ranged from 11.1 to 31.0 °C. In the laboratory, air temperature ranged 17.5 to 21.1 °C. The standardised *T*_*s*_ and *T*_*b_field*_ temperatures were estimated for 20 °C of ambient laboratory (*T*_*room*_) and for 20 °C of the two field temperatures, *T*_*air*_ (air temperature at breast height) and *T*_*microhabitat*_ (microhabitat temperature) experienced by individual butterflies in the spot where the individuals were located before capture (see Kleckova et al., 2014). For the laboratory measurements, the estimate of *T*_*s*_ at ambient temperature 20°C was calculated from a generalised linear model *T*_*s*_∼*T*_*room*_*+species* fitted to T_s_ estimates for all individuals (see above). We used the *predict()* function in R to obtain the estimate of T_s_ at ambient temperature 20 °C for each species. For the field observations, we again used the *predict()* function to infer *T*_*b_field*_ at ambient temperature 20 °C for each species. The dependence of body temperature *T*_*b_field*_ on both the ambient air (*T*_*a_field*_) and the microhabitat (*T*_*m_field*_) temperature was obtained from a previous study (Kleckova et al., 2014). Finally, we tested the relationship between the standardised *T*_*s*_ in the laboratory and the standardised *T*_*b_field*_ reached in the field for individual species by fitting a generalised linear model with Gaussian error distribution.

## 3. Results

### 3.1. Heating curves and species effect

Butterflies typically rested with closed wings after we positioned them in the experimental arena. Then, they usually opened their wings to a variable degree. Some individuals were closing and opening their wings repeatedly during the experiments. The wings heated up first (Figure 1), then, the temperature of the thorax gradually increased. Only a minority (12 individuals) of butterflies took off and flew away from the experimental arena during the measurements.

The increase of body temperature *T*_*b*_ in the studied *Erebia* butterflies was better described by the modified asymptotic exponential model (AIC = 583.8), which includes the heating rate parameter *b*, compared to the modified Michaelis-Menten model (AIC = 1051.3) (Table 2). The modified asymptotic exponential model predicted that the asymptotic temperature *T*_*s*_ is slightly more rapidly reached compared to the modified Michaelis-Menten model (Figure 2 and Table 2). The variability of the individual heating curves was best explained by a model including the random effect of individual identity nested within species (Table 2). However, differences in the shape of the heating curves among species were modest (Figure 3).

**Figure 3.**
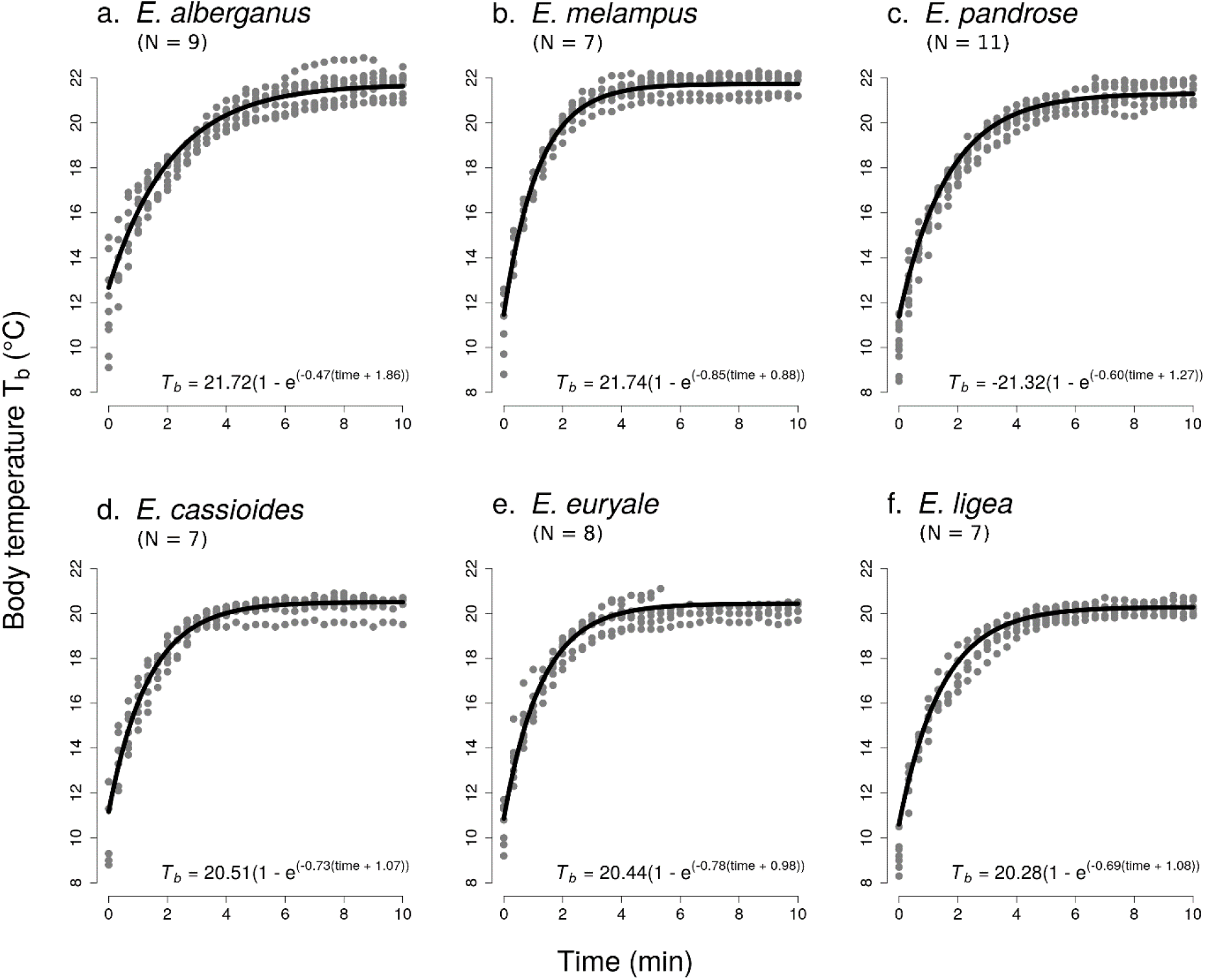
Heating of six *Erebia* butterfly species (49 individuals in total) under artificial source of heat and radiation under laboratory conditions. The lines display the fit of the modified asymptotic exponential model. The species-specific parameters of the model were estimated by a nonlinear mixed effects model with individual and species identity included as nested random effects. The parameter *b* describes the heating rate, i.e. the increase of the body temperature (*T*_*b*_) in *time* until it reaches the asymptotic body temperature (*T*_*s*_). The parameter *c* is the estimate of how long it would take to increase the body temperature from *T*_*b*_=0 °C to the real initial *T*_*b*_. Equations describing the fitted curves with species-specific parameter estimates are added.

### 3.2. Body weight and wing loading effects

The six species differed in their body weight (F_5,42_ = 18.13, P < 10^−8^) and wing loading (F_5,42_ = 13.51, P < 10^−7^, GLM with Gamma error distribution and log link function in both cases), although there was considerable intraspecific variation (Table 1). As expected, wing loading and body weight were strongly positively correlated (Pearson’s correlation coefficient = 0.84, t_46_ = 10.37, P <10^−12^). The majority of individuals of all species used in the experiments were males, except *E. pandrose* for which we measured 4 males and 7 females (Table 1).

Heating at the individual level was affected by both measures of body size. The heating rate *b* decreased with increasing body mass (GLMM, F = 7.17, P = 0.0105) and wing loading (GLMM, F = 4.40, P = 0.0419) (Figure 4). On the other hand, the asymptotic temperature *T*_*s*_ was independent of body mass (GLMM, F = 0.05, P = 0.8209) and wing loading (GLMM, F = 0.16, P = 0.6877). The results, thus, demonstrated that larger butterflies heated up more slowly but reached the same asymptotic temperature as smaller butterflies. The asymptotic body temperatures were on average 1.58 °C higher than the ambient air temperature (paired t-test, t_48_ = 13.394, P < 10^−15^). The GLMMs also included the effect of air temperature and humidity measured at the time of each observation. While *T*_*s*_ increased with the increasing room temperature (F = 46.53, P= <0.0001) and humidity (F = 5.12, P = 0.0294), the heating rate increased with the increasing room temperature (F = 13.82, P = 0.0011) and was not significantly affected by air humidity (F = 3.68, P = 0.0612).

**Figure 4.**
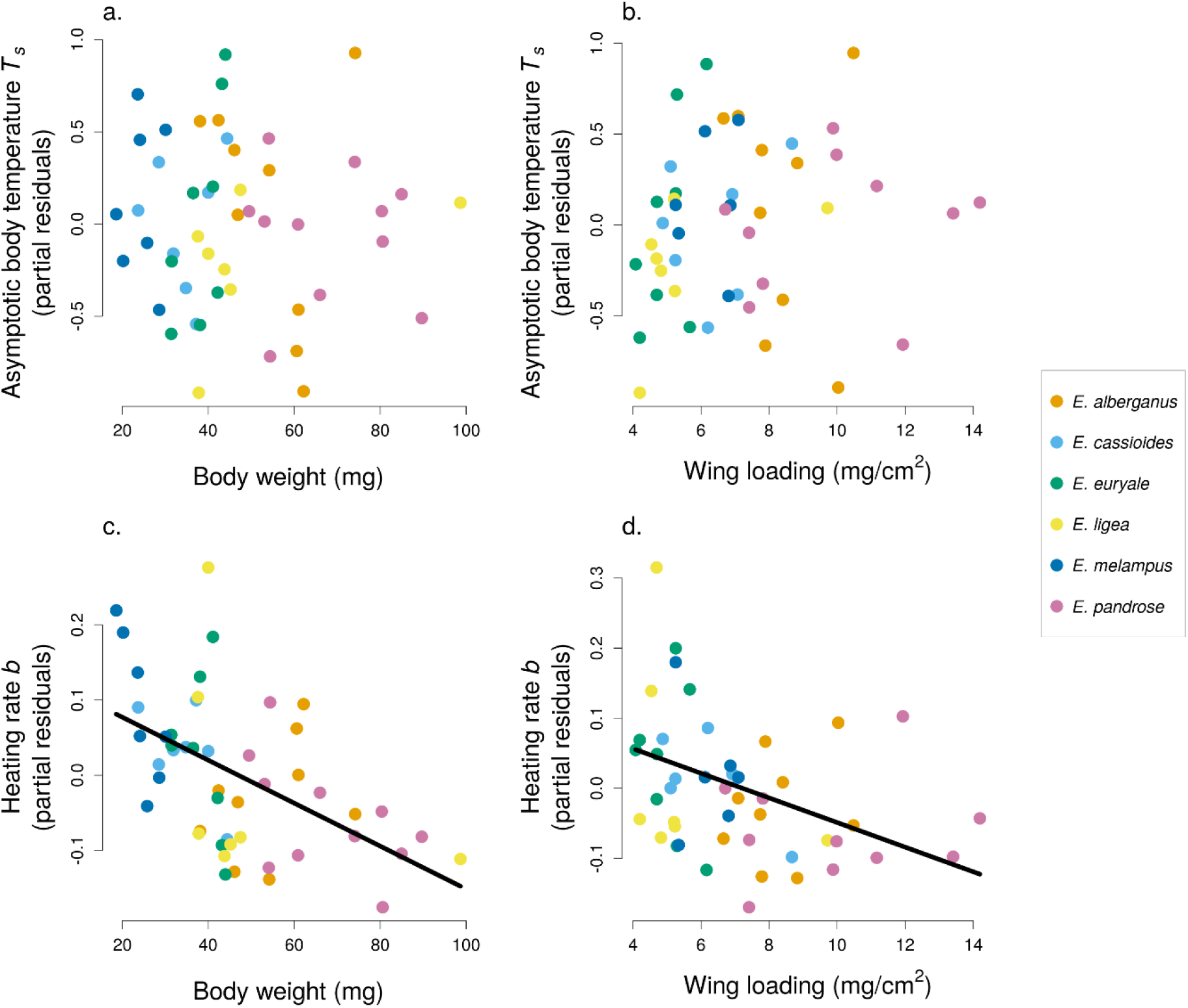
The effect of body weight and wing loading on the asymptotic body temperature (*T*_*s*_) and on the heating rate (*b*) of 49 individuals of six *Erebia* species. The *T*_*s*_ and *b* parameters describing heating of individuals under laboratory conditions were inferred from the modified asymptotic exponential models (see Figure 3). The y-axis shows partial residuals from a model which also included the room temperature and humidity at the time of each measurement. Each dot represents one individual, grouped into species according to colour.

### 3.3. Heating in the laboratory vs. body temperature in the field

The standardised asymptotic temperatures *T*_*s*_ measured in the laboratory (Figure 5) were not correlated with the standardised body temperature reached in field (*T*_*b_field*_). There was no statistically significant relationship regardless of whether *T*_*b_field*_ was standardised for air temperature *T*_*a_field*_ (GLM, F = 0.41, P = 0.5580) or for microhabitat temperature *T*_*m_field*_ (F = 0.84, P = 0.4107). It is noteworthy that the two species preferring rocky habitats, *E. cassioides* and *E. pandrose*, kept the highest body temperatures in the field despite their intermediate asymptotic temperatures measured in the laboratory (Figure 5). Also, the subalpine species *E. alberganus* and *E. melampus* kept higher body temperatures than the woodland species *E. euryale* and *E. ligea* measured both in the field and in the laboratory.

**Figure 5.**
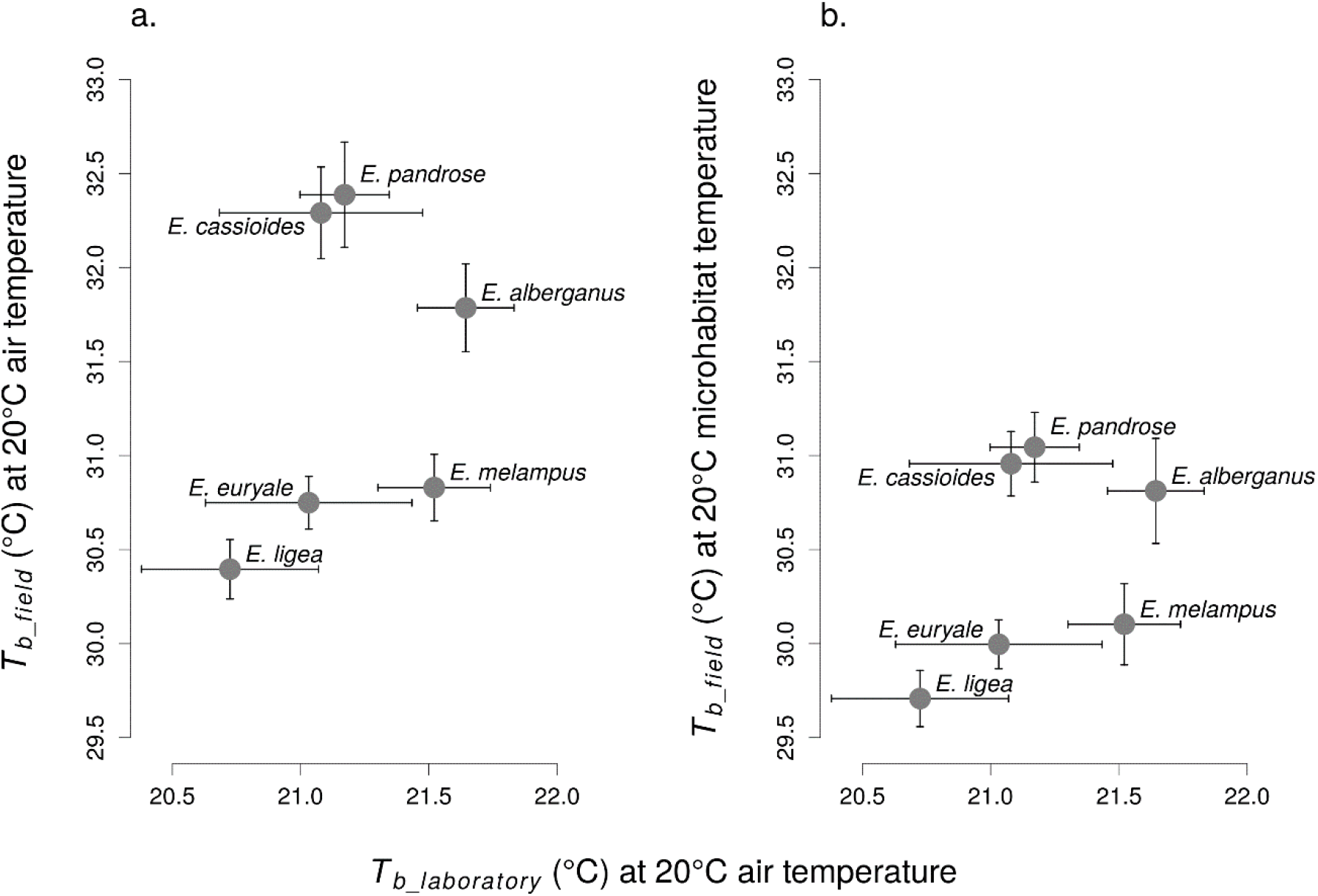
The correlation between *Erebia* butterfly species body temperatures reached under laboratory conditions (*T*_*b_laboratory*_) and mean body temperatures *T*_*b_field*_ measured in the field at 20°C of ambient air (panel a.) and microhabitat (panel b.) temperature. See the Methods section for more details. Estimates of *T*_*b*_ values are accompanied by standard errors.

## 4. Discussion

### 4.1. Species and individual effects on heating under laboratory conditions

The species-specific differences in body temperature heating in laboratory conditions were modest (Figure 3), but still detectable (Table 2). The better fit of the individual heating curves by models including the random effect of individual nested in species (Table 2) indicates that there is a noticeable effect of individual-level traits (e.g., species-specific physiological responses) in heating. However, the observed higher heating rate *b* in individuals with higher body weight and wing-loading regardless of species identity (Figure 4) points toward a major role of morphological traits. In other words, larger butterflies heated up more slowly than small species. This is in agreement with Konvicka et al. (2002), who documented that a large species in alpine grasslands, *Erebia euryale*, started activity at higher air temperatures and remained active longer under cloudy weather than a smaller sympatric species, *E. epiphron*. Our results are also in line with Berwaerts et al. (2001) and Kemp & Krockenberger (2004), who observed that larger individuals of nymphalid butterflies heated up more slowly than smaller ones. On the other hand, Neve & Hall (2016) found under controlled experimental conditions that smaller species of Mediterranean butterflies initiated flight at lower body temperatures than larger ones. In contrast, smaller *Erebia* species were in flight with higher body temperatures at low ambient temperatures on mountain slopes of European Alps (Kleckova et al., 2014). By studying the heating rates of *Erebia* species using a thermal camera in the laboratory, we confirmed that small body size allows faster heating and likely also facilitates active search for warmer microclimates during cold mornings or periods of adverse weather.

We have not confirmed our original expectation that larger *Erebia* species and individuals would accumulate higher body temperatures (higher asymptotic temperatures *T*_*s*_) in the laboratory, which was previously recorded in the butterfly species *Polyommatus icarus* (Lepidoptera: Lycaenidae) (Keyser et al., 2015). However, the similar asymptotic temperatures reached in our study might be partly explained by the low light intensity used in our experimental setup 45 W/m^2^), while solar flux is around 1000 W/m^2^ (Sterhov & Loshkarev, 2019). Future experiments should consider the importance of sufficient light intensity.

Other traits not included in our study, such as wing iridescence (Miaoulis & Heilman, 1998), emissivity, reflectance and absorbance (Lou et al., 2021; Schmitz, 1994) or fur (modified scales) thickness (Kingsolver & Moffat, 1982) on the body, may affect heating rates and body temperatures of butterflies. *Erebia* butterflies are iridescent with metallic reflections in freshly hatched butterflies, which may increase absorbance of solar influx (Bosi et al., 2008) in individuals with lower wing-wear. The effect of the density of the thoracic fur has been demonstrated, e.g., in *Colias* sulphur butterflies, where species inhabiting cold environments have dense fur on the thorax as it insulates against convective cooling (Watt, 1997). In the laboratory conditions, a small-bodied species, *E. melampus*, warmed up faster than the remaining species (Fig. 3), unlike the other small species, *E. cassiodes*. In the studied *Erebia* species, *E. cassioides* has the most prominent fur on the thorax. We assume that although the fur may provide better insulation in cold conditions (Kingsolver, 1988), it may also slow down heating.

The uniform heating rates in the several *Erebia* species in the laboratory conditions with limited possibilities of microclimate choice are rather expected based on the uniform morphology and behaviour under the heating source. Although we measured the heating rates in only approximately ten individuals (Table 1), this main result is robust. The technical restriction limiting the number of individuals was the availability of a single thermal camera and temperature-controlled room, which determined the number of individuals we were able to expose to experimental conditions after collecting them at a field site in Austria, which was remote from the lab (Czech Republic).

### 4.2. Habitat use

A previous study on adult microhabitat choice in the field demonstrated that woodland *Erebia* species kept lower body temperatures compared to species of open and rocky habitats (Kleckova et al., 2014). In congruence, the smallest difference in asymptotic temperature *T*_*s*_ reached in the laboratory and *T*_*b_field*_ under field conditions was found for the woodland species *E. ligea*. This indicates low buffering ability of this woodland species. Similarly, small differences in body temperatures measured in the field and in the laboratory were detected in another butterfly associated with shrubby grassland habitats and open woodlands, *Hamaeris lucina* (Riodinidae) (Bladon et al., 2020; Hayes et al., 2019). In contrast, the greatest difference between *T*_*s*_ and *T*_*b_field*_ was observed for species inhabiting rocky habitats and using stones or gravel as basking sites, *E. cassioides* and *E. pandrose*, indicating a likely active alteration of their body temperature in the wild by microclimate choice.

### 4.3. Heating curves were better described by the modified asymptotic exponential model

There is no consensus in the literature about the most appropriate model for fitting the heating curves describing the increase of body temperature *T*_*b*_ in time in insects. Exponential models were used to describe ectotherm heating, for example, to compare heating of colour morphs in the chameleon grasshopper (Umbers et al., 2013) and to infer the increase of internal body temperature in dung-beetles (Amore et al., 2017). Another model was the Michaelis-Menten equation, used e.g. to describe heating curves of a sea snail species by Muñoz et al. (2005). Complex models based on detailed physiological measurements of heat transfer within animal bodies and between them and their environment have been developed (Dreisig, 1984; Dzialowski & O’Connor, 2001), but they are impractical in ecological studies where detailed physiological parameters are difficult to obtain. In *Erebia* butterflies, the modified asymptotic exponential model explained the heating of body temperature better than the modified Michaelis-Menten model (Table 2). The main difference between the two models is that the exponential model allows for a sharper approach to the asymptote, while the Michaelis-Menten model is restricted to more gradual approach to the asymptote (see Fig. 2 for an illustrative example). Heating of the butterflies in our experiments was characterised by a near-linear initial increase of body temperature followed by a quick stabilisation of the body temperature at the asymptotic value (Fig. 3), which the exponential model fits better.

## 5. Implications

Studying the interactions of insect physical traits and their behavioural responses to environmental factors is important to understand and predict the impacts of climate change on community composition. When active thermoregulation by microhabitat choice was excluded under laboratory conditions, six closely related *Erebia* species differed minimally in passive heating, as measured by heating rate curves (Figure 3). We only detected a negative effect of individual body size on the rate of heating, but no effect on the asymptotic body temperature. Thus, the different thermal conditions and habitats found along elevational gradients have not driven divergent physiological constraints among species as we hypothesised, at least at the adult life stage. But, in the field, when adult butterflies have the opportunity to actively seek out suitable microclimates, the study species differed significantly in their body temperatures (Kleckova et al., 2014). This points toward a more prominent role of behavioural thermoregulation in these closely related butterflies than the physical traits (body size). However, further work is needed to test behavioural thermoregulation, e.g., by measuring active microclimate choice on a thermal gradient. The key role of behaviour in thermoregulation has been shown in a territorial butterfly (Rutowski et al., 1994) and in butterflies exposed to extreme temperatures (Tsai et al., 2020). The potential importance of behavioural thermoregulation implicated from this study suggests that the adult life stage of *Erebia* might not be under direct thermal selection, in agreement with Buckley et al. (2015). Thus, under increased anthropogenic pressure on *Erebia* populations, it seems necessary to safeguard a variety of microhabitats which can facilitate efficient behavioural thermoregulation of adult butterflies and increased survival of less mobile developmental stages (Abarca et al., 2019). This is in line with previous calls for management of mountain habitats supporting environmental heterogeneity (Habel et al., 2022) to increase species survival under ongoing climatic and other human-induced environmental changes (Lawson et al. 2014b).

## 6. Conclusions

Heating rates of adult *Erebia* butterflies under laboratory conditions differed slightly among species. However, smaller individuals heated up more quickly than larger ones regardless of species identity. This finding implies that small body size likely facilitates earlier initiation of butterfly daily activities (such as searching for warm microclimates) under unfavourable weather conditions. This is in an agreement with the observed higher body temperatures of small butterflies in the field on the mountain slopes of the Alps (Kleckova et al. 2014). However, the mild inter-specific differences in heating rates contrast with significant inter-specific differences in body temperatures documented in the field (Kleckova et al. 2014). To explain this finding, we speculate that microhabitat choice is a primary thermoregulatory mechanism. Thus, less mobile stages of butterfly developmental cycle are probably more severely affected by ongoing climatic change, which affects population trends (Konvicka et al., 2021) and species geographical distributions (Wilson et al., 2007).

## Acknowledgements

We are grateful to J. Skála and M. Česánek for recommending the study site, A. Sučhacková-Bartoňová for help with the thermal camera work, O. Nedvěd for providing practical advice, O. Terland and M. Konvička for the discussion of our results, E. Turner and one anonymous reviewer for thoughtful comments on a previous version of the manuscript, and to Marie Hronková for lending us the thermal camera. Funding was provided by the Czech Science Foundation (GAČR grant: GJ20-18566Y).

## Data availability

Data are accessible at https://doi.org/10.6084/m9.figshare.21836103. Full citation for data repository:

Kleckova I., Okrouhlik J., Svozil T., Matos-Maraví P. & Klecka J. (2022)

Body size, not species identity, drives body heating in alpine *Erebia* butterflies. Figshare Digital Repository Available at https://doi.org/10.6084/m9.figshare.21836103.

## References

Abarca, M., Larsen, E. A., & Ries, L. (2019). Heatwaves and Novel Host Consumption Increase Overwinter Mortality of an Imperiled Wetland Butterfly. Frontiers in Ecology and Evolution, 7. https://www.frontiersin.org/articles/10.3389/fevo.2019.00193

Abram, P. K., Boivin, G., Moiroux, J., & Brodeur, J. (2017). Behavioural effects of temperature on ectothermic animals: Unifying thermal physiology and behavioural plasticity. Biological Reviews of the Cambridge Philosophical Society, 92(4), 1859– 1876. https://doi.org/10.1111/brv.12312

Advani, N. K., Parmesan, C., & Singer, M. C. (2019). Takeoff temperatures in Melitaea cinxia butterflies from latitudinal and elevational range limits: A potential adaptation to solar irradiance. Ecological Entomology, 44(3), 389–396. https://doi.org/10.1111/een.12714

Amore, V., Hernández, M. I. M., Carrascal, L. M., & Lobo, J. M. (2017). Exoskeleton may influence the internal body temperatures of Neotropical dung beetles (Col. Scarabaeinae). PeerJ, 5, e3349. https://doi.org/10.7717/peerj.3349

Bates, D., Mächler, M., Bolker, B., & Walker, S. (2015). Fitting Linear Mixed-Effects Models Using lme4. Journal of Statistical Software, 67, 1–48. https://doi.org/10.18637/jss.v067.i01

Berwaerts, K., Van Dyck, H., Vints, E., & Matthysen, E. (2001). Effect of manipulated wing characteristics and basking posture on thermal properties of the butterfly Pararge aegeria (L.). Journal of Zoology, 255(2), 261–267. https://doi.org/10.1017/S0952836901001327

Bladon, A. J., Lewis, M., Bladon, E. K., Buckton, S. J., Corbett, S., Ewing, S. R., Hayes, M. P., Hitchcock, G. E., Knock, R., Lucas, C., McVeigh, A., Menéndez, R., Walker, J. M., Fayle, T. M., & Turner, E. C. (2020). How butterflies keep their cool: Physical and ecological traits influence thermoregulatory ability and population trends. Journal of Animal Ecology, 89(11), 2440–2450. https://doi.org/10.1111/1365-2656.13319

Bosi, S. G., Hayes, J., Large, M. C. J., & Poladian, L. (2008). Color, iridescence, and thermoregulation in Lepidoptera. Applied Optics, 47(29), 5235–5241. https://doi.org/10.1364/AO.47.005235

Buckley, L. B., Ehrenberger, J. C., & Angilletta Jr, M. J. (2015). Thermoregulatory behaviour limits local adaptation of thermal niches and confers sensitivity to climate change. Functional Ecology, 29(8), 1038–1047. https://doi.org/10.1111/1365-2435.12406

Cizek, O., Bakesová, A., Kuras, T., Benes, J., & Konvicka, M. (2003). Vacant niche in alpine habitat: The case of an introduced population of the butterfly Erebia epiphron in the Krkonoše Mountains. Acta Oecologica, 24(1), 15–23. https://doi.org/10.1016/S1146-609X(02)00004-8

Cros, S., Cerdá, X., & Retana, J. (1997). Spatial and temporal variations in the activity patterns of Mediterranean ant communities. Écoscience, 4(3), 269–278. https://doi.org/10.1080/11956860.1997.11682405

Dreisig, H. (1984). Control of body temperature in shuttling ectotherms. Journal of Thermal Biology, 9(4), 229–233. https://doi.org/10.1016/0306-4565(84)90001-9

Dreisig, H. (1995). Thermoregulation and flight activity in territorial male graylings, Hipparchia semele (Satyridae), and large skippers, Ochlodes venata (Hesperiidae). Oecologia, 101(2), 169–176. https://doi.org/10.1007/BF00317280

Dzialowski, E. M., & O’Connor, M. P. (2001). Thermal time constant estimation in warming and cooling ectotherms. Journal of Thermal Biology, 26(3), 231–245. https://doi.org/10.1016/s0306-4565(00)00050-4

Forsman, A. (2000). Some like it hot: Intra-Population Variation in behavioral Thermoregulation in Color-Polymorphic pygmy Grasshoppers. Evolutionary Ecology, 14(1), 25–38. https://doi.org/10.1023/A:1011024320725

Gallego, B., Verdú, J. R., & Lobo, J. M. (2018). Comparative thermoregulation between different species of dung beetles (Coleoptera: Geotrupinae). Journal of Thermal Biology, 74, 84–91. https://doi.org/10.1016/j.jtherbio.2018.03.009

Habel, J. C., Schmitt, T., Gros, P., & Ulrich, W. (2022). Breakpoints in butterfly decline in Central Europe over the last century. Science of The Total Environment, 851, 158315. https://doi.org/10.1016/j.scitotenv.2022.158315

Hayes, M. P., Hitchcock, G. E., Knock, R. I., Lucas, C. B. H., & Turner, E. C. (2019). Temperature and territoriality in the Duke of Burgundy butterfly, Hamearis lucina. Journal of Insect Conservation, 23(4), 739–750. https://doi.org/10.1007/s10841-019-00166-6

Heinrich, B. (1986). Thermoregulation and Flight Activity Satyrine, Coenonympha Inornata (Lepidoptera: Satyridae). Ecology, 67(3), 594–597. https://doi.org/10.2307/1937682

Heinrich, B. (1995). Insect thermoregulation. Endeavour, 19(1), 28–33. https://doi.org/10.1016/0160-9327(95)98891-I

Kemp, D. J., & Krockenberger, A. K. (2002). A novel method of behavioural thermoregulation in butterflies. Journal of Evolutionary Biology, 15(6), 922–929. https://doi.org/10.1046/j.1420-9101.2002.00470.x

Kemp, D. J., & Krockenberger, A. K. (2004). Behavioural thermoregulation in butterflies: The interacting effects of body size and basking posture in Hypolimnas bolina (L.) (Lepidoptera : Nymphalidae). Australian Journal of Zoology, 52(3), 229–239. https://doi.org/10.1071/zo03043

Keyser, R. D., Breuker, C. J., Hails, R. S., Dennis, R. L. H., & Shreeve, T. G. (2015). Why Small Is Beautiful: Wing Colour Is Free from Thermoregulatory Constraint in the Small Lycaenid Butterfly, Polyommatus icarus. PLOS ONE, 10(4), e0122623. https://doi.org/10.1371/journal.pone.0122623

Kingsolver, J. G. (1983). Thermoregulation and Flight in Colias Butterflies: Elevational Patterns and Mechanistic Limitations. Ecology, 64(3), 534–545. https://doi.org/10.2307/1939973

Kingsolver, J. G. (1985a). Thermoregulatory significance of wing melanization in Pieris butterflies (Lepidoptera: Pieridae): physics, posture, and pattern. Oecologia, 66(4), 546–553. https://doi.org/10.1007/BF00379348

Kingsolver, J. G. (1985b). Thermoregulatory significance of wing melanization in Pieris butterflies (Lepidoptera: Pieridae): physics, posture, and pattern. Oecologia, 66(4), 546–553. https://doi.org/10.1007/BF00379348

Kingsolver, J. G. (1988). Thermoregulation, Flight, and the Evolution of Wing Pattern in Pierid Butterflies: The Topography of Adaptive Landscapes. American Zoologist, 28(3), 899–912. https://doi.org/10.1093/icb/28.3.899

Kingsolver, J. G., & Moffat, R. J. (1982). Thermoregulation and the determinants of heat transfer in Colias butterflies. Oecologia, 53(1), 27–33. https://doi.org/10.1007/BF00377132

Kirkpatrick, W. H., & Sheldon, K. S. (2022). Experimental increases in temperature mean and variance alter reproductive behaviours in the dung beetle Phanaeus vindex. Biology Letters, 18(7), 20220109. https://doi.org/10.1098/rsbl.2022.0109

Kleckova, I., & Klecka, J. (2016). Facing the Heat: Thermoregulation and Behaviour of Lowland Species of a Cold-Dwelling Butterfly Genus, Erebia. PLOS ONE, 11(3), e0150393. https://doi.org/10.1371/journal.pone.0150393

Kleckova, I., Konvicka, M., & Klecka, J. (2014). Thermoregulation and microhabitat use in mountain butterflies of the genus Erebia: Importance of fine-scale habitat heterogeneity. Journal of Thermal Biology, 41, 50–58. https://doi.org/10.1016/j.jtherbio.2014.02.002

Konvicka, M., Beneš, J., & Kuras, T. (2002). Microdistribution and diurnal behaviour of two sympatric mountainous butterflies (Erebia epiphron and E. euryale): Relations to vegetation and weather. Biologia, 57, 225–235.

Konvicka, M., Kuras, T., Liparova, J., Slezak, V., Horázná, D., Klečka, J., & Kleckova, I. (2021). Low winter precipitation, but not warm autumns and springs, threatens mountain butterflies in middle-high mountains. PeerJ, 9, e12021. https://doi.org/10.7717/peerj.12021

Konvicka, M., Maradova, M., Benes, J., Fric, Z., & Kepka, P. (2003). Uphill shifts in distribution of butterflies in the Czech Republic: Effects of changing climate detected on a regional scale. Global Ecology and Biogeography, 12(5), 403–410. https://doi.org/10.1046/j.1466-822X.2003.00053.x

Kuras, T., Benes, J., Fric, Z., & Konvicka, M. (2003). Dispersal patterns of endemic alpine butterflies with contrasting population structures: Erebia epiphron and E. sudetica. Population Ecology, 45(2), 115–123. https://doi.org/10.1007/s10144-003-0144-x

Kuznetsova, A., Brockhoff, P. B., & Christensen, R. H. B. (2017). lmerTest Package: Tests in Linear Mixed Effects Models. Journal of Statistical Software, 82, 1–26. https://doi.org/10.18637/jss.v082.i13

Lawson, C. R., Bennie, J., Hodgson, J. A., Thomas, C. D., & Wilson, R. J. (2014). Topographic microclimates drive microhabitat associations at the range margin of a butterfly. Ecography, 37(8), 732–740. https://doi.org/10.1111/ecog.00535

Lawson, C. R., Bennie, J. J., Thomas, C. D., Hodgson, J. A., & Wilson, R. J. (2014). Active Management of Protected Areas Enhances Metapopulation Expansion Under Climate Change. Conservation Letters, 7(2), 111–118. https://doi.org/10.1111/conl.12036

Lou, C., An, S., Yang, R., Zhu, H., Shen, Q., Jiang, M., Fu, B., Tao, P., Song, C., Deng, T., & Shang, W. (2021). Enhancement of infrared emissivity by the hierarchical microstructures from the wing scales of butterfly Rapala dioetas. APL Photonics, 6(3), 036101. https://doi.org/10.1063/5.0039079

MacLean, H. J., Higgins, J. K., Buckley, L. B., & Kingsolver, J. G. (2016). Morphological and physiological determinants of local adaptation to climate in Rocky Mountain butterflies. Conservation Physiology, 4(1), cow035. https://doi.org/10.1093/conphys/cow035

Mattila, A. L. K. (2015). Thermal biology of flight in a butterfly: Genotype, flight metabolism, and environmental conditions. Ecology and Evolution, 5(23), 5539–5551. https://doi.org/10.1002/ece3.1758

Miaoulis, I. N., & Heilman, B. D. (1998). Butterfly Thin Films Serve as Solar Collectors. Annals of the Entomological Society of America, 91(1), 122–127. https://doi.org/10.1093/aesa/91.1.122

Muñoz, J. L. P., Randall Finke, G., Camus, P. A., & Bozinovic, F. (2005). Thermoregulatory behavior, heat gain and thermal tolerance in the periwinkle Echinolittorina peruviana in central Chile. Comparative Biochemistry and Physiology Part A: Molecular & Integrative Physiology, 142(1), 92–98. https://doi.org/10.1016/j.cbpa.2005.08.002

Nève, G., & Després, L. (2020). Cold adaptation across the elevation gradient in an alpine butterfly species complex. Ecological Entomology, 45(5), 997–1003. https://doi.org/10.1111/een.12875

Neve, G., & Hall, C. (2016). Variation of thorax flight temperature among twenty Australian butterflies (Lepidoptera: Papilionidae, Nymphalidae, Pieridae, Hesperiidae, Lycaenidae). European Journal of Entomology, 113, 571–578. https://doi.org/10.14411/eje.2016.077

Pinheiro, J., Bates, D., & R Core Team. (2022). Nlme: Linear and Nonlinear Mixed Effects Models. R package version 3.1-157, https://CRAN.R-project.org/package=nlme.

Pivnick, K. A., & McNeil, J. N. (1986). Sexual Differences in the Thermoregulation of Thymelicus Lineola Adults (Lepidoptera: Hesperiidae). Ecology, 67(4), 1024–1035. https://doi.org/10.2307/1939825

Polic, D., Fiedler, K., Nell, C., & Grill, A. (2014). Mobility of ringlet butterflies in high-elevation alpine grassland: Effects of habitat barriers, resources and age. Journal of Insect Conservation, 18(6), 1153–1161. https://doi.org/10.1007/s10841-014-9726-5

R Core Team. (2022). R: A language and environment for statistical computing. R Foundation for Statistical Computing, Vienna, Austria. https://www.R-project.org/.

Radchuk, V., Turlure, C., & Schtickzelle, N. (2013). Each life stage matters: The importance of assessing the response to climate change over the complete life cycle in butterflies. Journal of Animal Ecology, 82(1), 275–285. https://doi.org/10.1111/j.1365-2656.2012.02029.x

Rasband, W. S. (1997). ImageJ, U. S. National Institutes of Health, Bethesda, Maryland, USA (v.1.46). https://imagej.nih.gov/ij/

Rödder, D., Schmitt, T., Gros, P., Ulrich, W., & Habel, J. C. (2021). Climate change drives mountain butterflies towards the summits. Scientific Reports, 11(1), 14382. https://doi.org/10.1038/s41598-021-93826-0

Rutowski, R. L., Demlong, M. J., & Leffingwell, T. (1994). Behavioural thermoregulation at mate encounter sites by male butterflies (Asterocampa leilia, Nymphalidae). Animal Behaviour, 48(4), 833–841. https://doi.org/10.1006/anbe.1994.1307

Schmitz, H. (1994). Thermal characterization of butterfly wings—1. Absorption in relation to different color, surface structure and basking type. Journal of Thermal Biology, 19(6), 403–412. https://doi.org/10.1016/0306-4565(94)90039-6

Shreeve, T. G. (1984). Habitat Selection, Mate Location, and Microclimatic Constraints on the Activity of the Speckled Wood Butterfly Pararge Aegeria. Oikos, 42(3), 371–377. https://doi.org/10.2307/3544407

Sistri, G., Menchetti, M., Santini, L., Pasquali, L., Sapienti, S., Cini, A., Platania, L., Balletto, E., Barbero, F., Bonelli, S., Casacci, L. P., Dincă, V., Vila, R., Mantoni, C., Fattorini, S., & Dapporto, L. (2022). The isolated Erebia pandrose Apennine population is genetically unique and endangered by climate change. Insect Conservation and Diversity, 15(1), 136–148. https://doi.org/10.1111/icad.12538

Slamova, I., Klecka, J., & Konvicka, M. (2011). Diurnal Behavior and Habitat Preferences of Erebia aethiops, an Aberrant Lowland Species of a Mountain Butterfly Clade. Journal of Insect Behavior, 24(3), 230–246. https://doi.org/10.1007/s10905-010-9250-8

Slamova, I., Klecka, J., & Konvicka, M. (2013). Woodland and grassland mosaic from a butterfly perspective: Habitat use by Erebia aethiops (Lepidoptera: Satyridae). Insect Conservation and Diversity, 6(3), 243–254. https://doi.org/10.1111/j.1752-4598.2012.00212.x

Sterhov, A. I., & Loshkarev, I. Y. (2019). Determination of the proportion of natural light in solar radiation using the method of conversion of lighting units into energy. Journal of Physics: Conference Series, 1353(1), 012002. https://doi.org/10.1088/1742-6596/1353/1/012002

Stuhldreher, G., & Fartmann, T. (2018). Threatened grassland butterflies as indicators of microclimatic niches along an elevational gradient – Implications for conservation in times of climate change. Ecological Indicators, 94, 83–98. https://doi.org/10.1016/j.ecolind.2018.06.043

Tsai, C.-C., Childers, R. A., Nan Shi, N., Ren, C., Pelaez, J. N., Bernard, G. D., Pierce, N. E., & Yu, N. (2020). Physical and behavioral adaptations to prevent overheating of the living wings of butterflies. Nature Communications, 11(1), Article 1. https://doi.org/10.1038/s41467-020-14408-8

Umbers, K. D. L., Herberstein, M. E., & Madin, J. S. (2013). Colour in insect thermoregulation: Empirical and theoretical tests in the colour-changing grasshopper, Kosciuscola tristis. Journal of Insect Physiology, 59(1), 81–90. https://doi.org/10.1016/j.jinsphys.2012.10.016

Verdu, J. R., Alba-Tercedor, J., & Jimenez-Manrique, M. (2012). Evidence of Different Thermoregulatory Mechanisms between Two Sympatric Scarabaeus Species Using Infrared Thermography and Micro-Computer Tomography | PLOS ONE. PLoS ONE, 7(3), | e3391.

Watt, W. B. (1997). Accuracy, Anecdotes, and Artifacts in the Study of Insect Thermal Ecology. Oikos, 80(2), 399. https://doi.org/10.2307/3546607

Wickman, P. O. (2009). Thermoregulation and habitat use in butterflies. In Settele, J., Shreeve, T., Konvicka, M., Van Dyck, H. (Eds.). Ecology of Butterflies in Europe (pp. 55–61).

Wickman, P.-O. (1988). Dynamics of mate-searching behaviour in a hilltopping butterfly, Lasiommata megera (L.): The effects of weather and male density. Zoological Journal of the Linnean Society, 93(4), 357–377. https://doi.org/10.1111/j.1096-3642.1988.tb01367.x

Willmer, P. G. (1982). Microclimate and the Environmental Physiology of Insects. In M. J. Berridge, J. E. Treherne, & V. B. Wigglesworth (Eds.), Advances in Insect Physiology (Vol. 16, pp. 1–57). Academic Press. https://doi.org/10.1016/S0065-2806(08)60151-4

Wilson, R. J., Bennie, J., Lawson, C. R., Pearson, D., Ortúzar-Ugarte, G., & Gutiérrez, D. (2015). Population turnover, habitat use and microclimate at the contracting range margin of a butterfly. Journal of Insect Conservation, 19(2), 205–216. https://doi.org/10.1007/s10841-014-9710-0

Wilson, R. J., Gutiérrez, D., Gutiérrez, J., & Monserrat, V. J. (2007). An elevational shift in butterfly species richness and composition accompanying recent climate change. Global Change Biology, 13(9), 1873–1887. https://doi.org/10.1111/j.1365-2486.2007.01418.x

